# Are fungi capacitors?

**DOI:** 10.1101/2023.05.28.542674

**Authors:** Konrad Szaciłowski, Alexander E. Beasley, Krzysztof Mech, Andrew Adamatzky

## Abstract

The emerging field of living technologies aims to create new functional hybrid materials in which living systems interface and interact with inanimate ones. Combining research into living technologies with emerging developments in computing architecture has enabled the generation of organic electronics from plants and slime mould. Here, we expand on this work by studying capacitive properties of a substrate colonised by mycelium of grey oyster fungi, *Pleurotus ostreatus*. Capacitors play a fundamental role in traditional analogue and digital electronic systems and have a range of uses including sensing, energy storage and filter circuits. Mycelium has the potential to be used as an organic replacement for traditional capacitor technology. Here, were show that the capacitance of mycelium is in the order of hundreds of picofarads and at the same time a voltage-dependent pseudocapacitance of the order of hundreds of microfarads. We also demonstrate that the charge density of the mycelium ‘dielectric’ decays rapidly with increasing distance from the source probes. This is important as it indicates that small cells of mycelium could be used as a charge carrier or storage medium, when employed as part of an array with reasonable density.

## 1 Introduction

The study of novel substrates for sensing, storing and processing information draws on work from the fields of unconventional computing, living technology and organic electronics. The field of unconventional computing aims to define the principles of information processing in living, physical and chemical systems and applies this knowledge to the development of future computing devices and architectures [1]. Research into living technologies is focused on the co-functional integration of animate and non-organic systems [2]. Finally, organic electronics [3, 4] looks to use naturally occurring materials as analogues to traditional semi-conductor circuits [5], which often requires functionalisation using polymers or metallic compounds to exploit ionic movement [6]. The development of organic electronics promises a technology that is low-cost and has low production temperature requirements [7]. However, difficulties such as relatively low gain of organic transistors (approx. 5), and the behavioural variability inherently means that there is a large amount of research effort being placed into organic thin film transistors [8, 9, 10, 11], organic LEDs [12, 13], and organic capacitors [14, 15, 16]. The capacitive properties of a device allows it to either store energy or react to AC/DC signals differently. There are a number of applications in which this property may be utilised, such as energy harvesting [17], memory [18], or filter circuits [19]. Hybrid electronic circuits are a concept that looks to combine traditional silicon, semi-conductor devices with elements found in nature [20, 21].

The capacitive properties of living tissues [22] have a wide range of potential applications, e.g. the estimation of a plan root system size [23, 24], quantifying the DNA content of eukaryotic cells [25], analysing water transport pathways in plants [26], measuring heat injury in plants [27], measuring contents of minerals in bones [28], gauging firmness of apples [29], sugar contents of citrus fruits [30], maturity of avocados [31], estimating depth of epidermal barriers [32], studies of endo- and exocytosis of single cells [33], and approximating mass and morphology of microbial colonies [34, 35, 36].

Among various possible biological substrates for electronics, fungal mycelia are one of the most promising materials. In [37] we proposed a novel field of bioelectronics — Fungal Electronics, which includes novel and original electronic devices made of living mycelium. At the same year other groups similarly proposed to utilise fungal mycelium skin for sustainable electronics [38].

Continuously increasing demand for electronics of new capabilities toward sensing, signal analysis, data acquisition, and information processing pushes scientists for searching for unconventional approaches creating new possibilities that may allow fulfilling the growing requirements of users. The utilisation of fungal mycelium seems to be an interesting approach that may enable the fabrication of a new class of green wearable electronics of the future.

In the present study we focus on the capacitive properties of the mycelium of the grey oyster fungi *Pleurotus ostreatus* for several reasons.

Firstly, research into the capacitive properties of fungi is lacking, despite their huge potential for bioelectronic applications. Fungi are the largest, most widely distributed and oldest group of living organisms on the planet [39]. The smallest fungi are microscopic single cells, the largest, *Armillaria bulbosa*, occupies 15 hectares and weighs 10 tons [40]. Fungi sense light, chemicals, gases, gravity and electric fields [41] as well as demonstrating mechanosensing behaviour [42]. Thus, their electrical properties can be tuned via various inputs.

Secondly, fungi have the potential to be used as distributed living computing devices, i.e. large-scale networks of mycelium, which collect and analyse information about environment and execute some decision making processes [43].

Finally, there is a growing interest in developing buildings from pre-fabricated blocks of substrates colonised by fungi [44, 45, 46]. A recent initiative aims to grow monolithic constructions in which living mycelium coexists with dried mycelium, functionalised with nanoparticles and polymers [47]. In such a case, fungi could be act as optical, tactile and chemical sensors, fuse and process information and perform decision making computations [43].

Providing local charge to areas of mycelium allows the storage of information inside the substrate. Identifying the area around which the induced charge can be detected allows the construction of an array in the substrate where each cell can contain individual bits of information. Determining the capacitive properties of fungi takes a step towards the realisation of fungal analogue circuits — circuits that use fungi to replace traditional semiconductors.

The rest of this paper is organised as follows. Section 2 describes the experimental methods used for the analysis of the substrate. Section 3 presents the results with discussions. Finally, conclusions are drawn in Sect. 4.

## 2 Experimental method

Mycelium of the grey oyster fungi *Pleurotus ostreatus* (Ann Miller’s Speciality Mushrooms Ltd, UK) was cultivated on damp wood shavings (Fig. 1(a)). Control samples of the growth medium, wood shavings, were not colonised by mycelium. Iridium-coated stainless steel sub-dermal needles with twisted cables (Spes Medica SRL, Italy) were inserted in the colonised substrate. An LCR meter (BK Precision, model 891) was used to provide a nominal reading of the capacitance of the sample with probes at 10 mm, 20 mm, 40 mm and 50 mm separation. Cyclic voltammograms in two electrode setup has been recorded with with SP-150 potentiostat (Bio-Logic, France) in the dark. Impedance spectra in the 1 Hz-100 kHz frequency range were recorded with Bode 100 vector network analyser (Omicron Labs, Austria). Voltage dependence of pseudocapacitance was measured using chronocoulometric module of Bio-Logic potentiostat.

**Figure 1:**
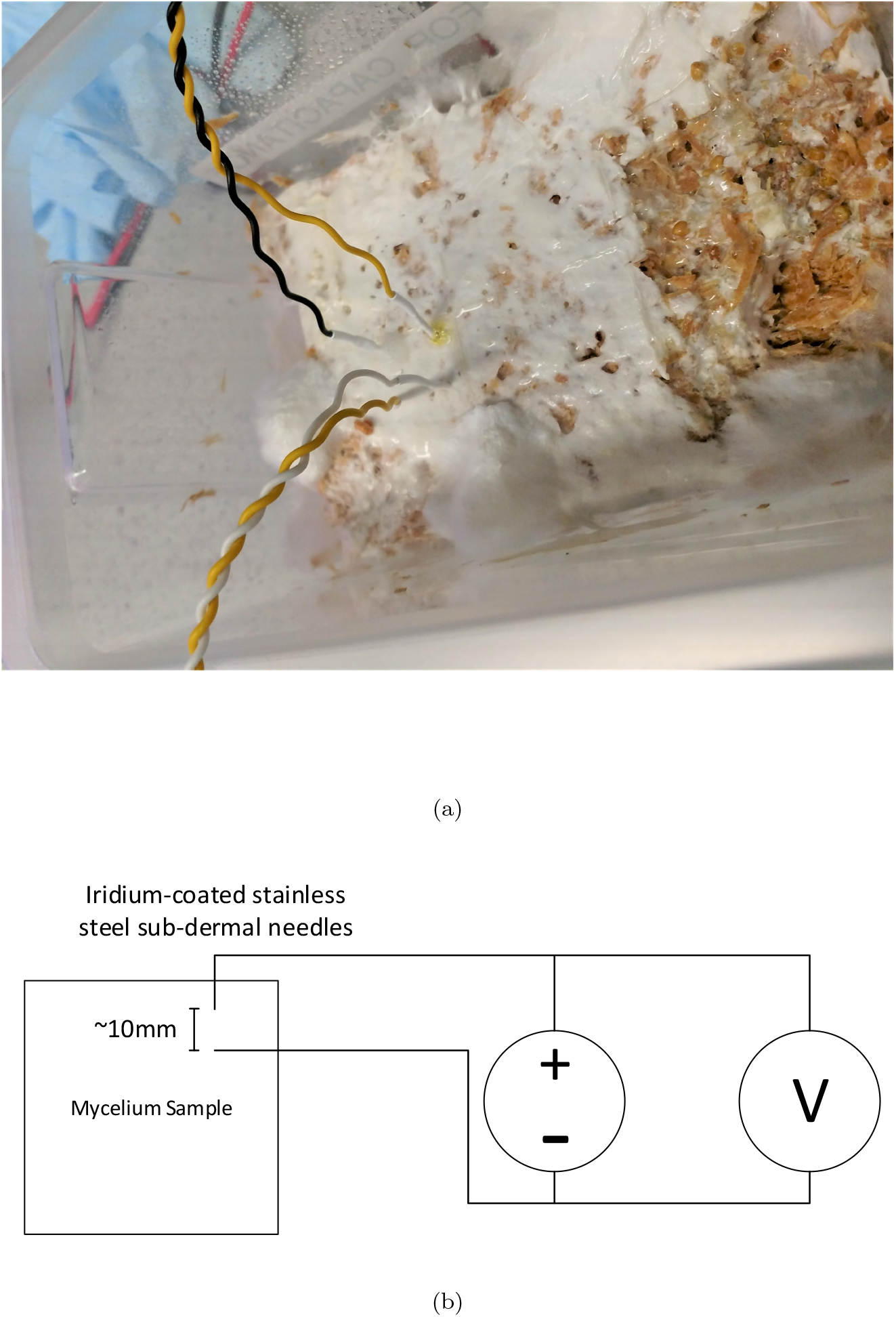
Experimental setup. (a) A sample of the mycelium under test. (b) Voltage discharge measuring set up for mycelium samples

The samples were charged using a bench top DC power supply (BK precision 9206) to 50 V. The power supply output was de-activated and the discharge curve was measured using a bench top digital multi-meter (DMM) Fluke 8846A (Fig. 1(b)). To fully characterise the capacitance of the samples, both the charge and discharge curves were monitored [48] with a number of probe separations (e.g. 10 mm, 20 mm, 40 mm, and 50 mm). Measurements from the DMM were automated through a serial terminal from a host PC. All bench-top equipment was high impedance to limit power lost through leakage in the test equipment. All plots were generated using MATLAB. The sample interval was approximately 0.33 s. Multiple samples and repeated experimental recordings were used to increase the statistical significance of the findings.

## 3 Results

### Capacitance measurement

Samples of the growth medium and mycelium were measured for their capacitance using a standard bench top LCR meter (Tab. 1). The measured capacitance value of the mycelium substrate was two to four fold greater than that of the growth medium alone. The value recorded for dry wood shavings corresponds to the capacitance of the twisted cable inserted into the sample and served as a background value. Increased humidity of the pristine substrate is higher. The values of capacitance were also effected by the moisture content such that, if the capacitance of the mycelium was measured straight after it is sprayed with water, the capacitance typically increased compared to that of dry substrate.

**Table 1:** Capacitance of mycelium and growth mediums.

| Sample | Electrode spacing | Capacitance (pF) |
| --- | --- | --- |
| Dry wood shavings | 10 mm | 37.6 |
|  | 20 mm | 37.4 |
| Damp wood shavings | 10 mm | 57 |
|  | 20 mm | 58.6 |
| Mycelium (drying) | 10 mm | 184 |
|  | 20 mm | 144 |
|  | 30 mm | 146 |
|  | 40 mm | 120 |
|  | 50 mm | 118 |
| Mycelium (freshly watered) | 10 mm | 193 |
|  | 20 mm | 186 |
|  | 30 mm | 134 |
|  | 40 mm | 125 |
|  | 50 mm | 139 |

### Discharge characteristics

Discharge characteristics of the growth medium and mycelium samples are shown in Figs. 2–3. Discharge curves were produced by setting up the DC power supply and DMM in parallel with each other. The substrate being tested is then charged to 50 V and the power supply output is disabled. The DMM continued to periodically measure the remaining charge in the substrate for a period of time. The sample interval was approximately 0.33 s.

**Figure 2:**
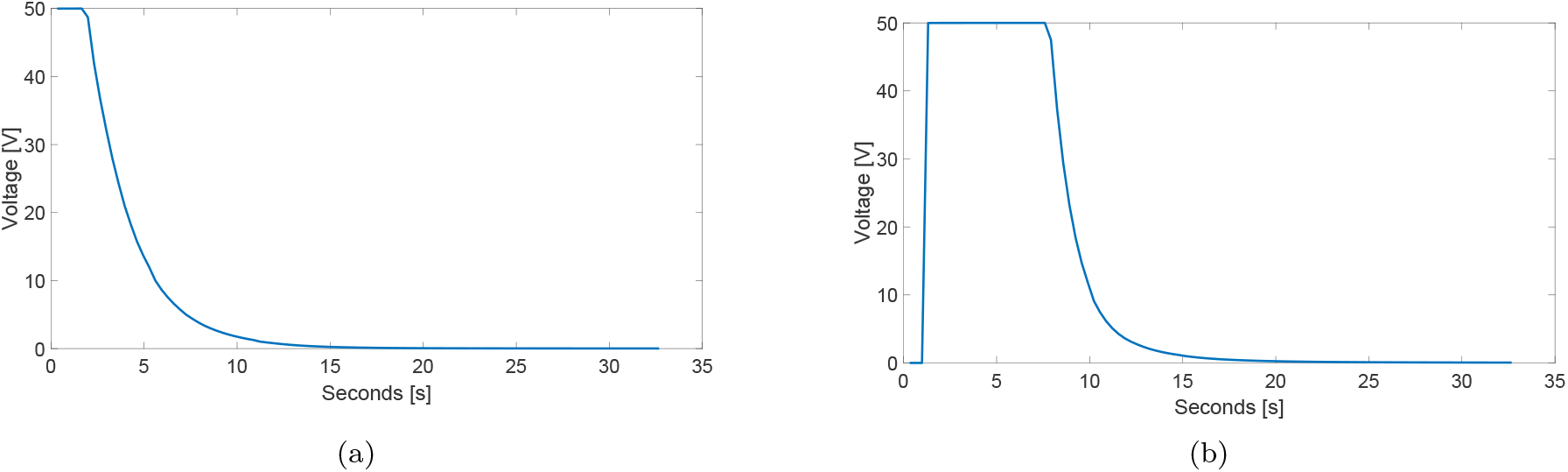
Discharge of a substrate after being charged to 50 V with probes separation of 10 mm. (a) Dry wood shavings. (b) Damp wood shavings — shavings are immersed in water for half and hour after which excess water is drained. Data are discrete. Line is for eye guidance only.

The discharge curves for both the growth medium and the mycelium are very steep - approximated by an exponential (1) .

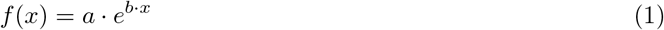

Where the parameters for 95% fitness for different mediums can be found in Table 2.

**Table 2:** Discharge curve fitness approximation coefficients (with 95% confidence bounds)

| Sample | $a$ | $b$ |
| --- | --- | --- |
| Dry wood shavings | 104.9 (102.6, 107.2) | -0.4079 (-0.4151, -0.4007) |
| Damp wood shavings | 6205 (5164, 7245) | -0.6261 (-0.6464, -0.6058) |
| Mycelium w/10 mm probe separation | 2276 (2198, 2354) | -0.4446 (-0.4481, -0.441) |
| Mycelium w/20 mm probe separation | 2688 (2544, 2833) | -0.4639 (-0.4695, -0.4583) |
| Mycelium w/40 mm probe separation | 1569 (1458, 1680) | -0.3948 (-0.4021, -0.3875) |
| Mycelium w/50 mm probe separation | 1413 (1311, 1516) | -0.3817 (-0.3891, -0.3743) |

The discharge time is governed by equation *τ* = *RC*, where *τ* is the time constant, *R* is a resistance, and *C* is a capacitance.

With capacitance in the order of pico-Farads, and input impedance of the source and measurement equipment in the order of mega-Ohms, it is expected that the discharge will be in the order of seconds.

Comparing the discharge curves of the growth medium to that of the mycelium samples, it is evident that the discharge was not as steep in the mycelium due to the increase in capacitance over the growth medium. However, it was still in the pico-Farad range and, therefore, the majority of charge was lost after just over 5 s. Increasing the separation distance of the probes (Fig. 3) had only a minimal effect on the capacitance of the substrate (shown in Tab. 1), and therefore minimal effect on the discharge curve.

**Figure 3:**
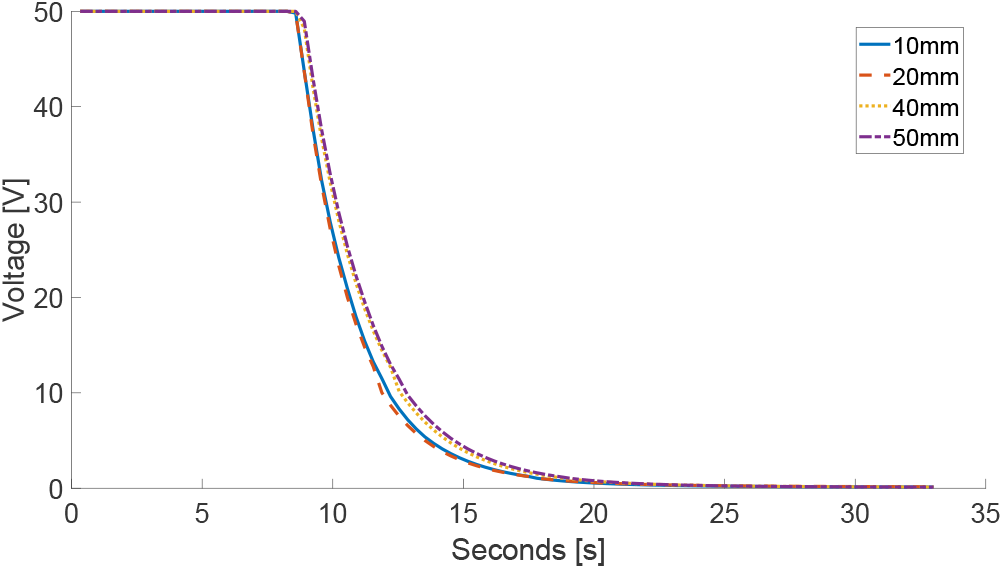
Mycelium sample is charged to 50 V then allowed to discharge. Electrode spacing is varied (10 mm, 20 mm, 40 mm and 50 mm). Data are discrete. Line is for eye guidance only.

Observing the charging and discharging behaviour of the sample around the source electrodes helps to build a better picture of the current density of the mycelium substrate. Figure 4 shows how we placed the measurement equipment electrodes 10 mm away from the source electrodes in the mycelium, in three different locations shown in Fig. 5. The sample was then charged for approximately 10 mins before the supply was turned off. The electrodes placed in ‘series’ (locations (a) and (b) on Fig. 5) with the charge electrodes (Fig. 4(a)) show minimum voltage detected beyond the supply electrodes in the horizontal plane, when the supply is active. Placing the measurement probes in ‘parallel’ (location (c) on Fig. 5) with the supply probes (Fig. 4(b)) demonstrates the fact that considerably more current is conducted between the supply probes in the vertical plane. When the supply is de-activated, the voltage around the supplies collapses almost instantly.

**Figure 4:**
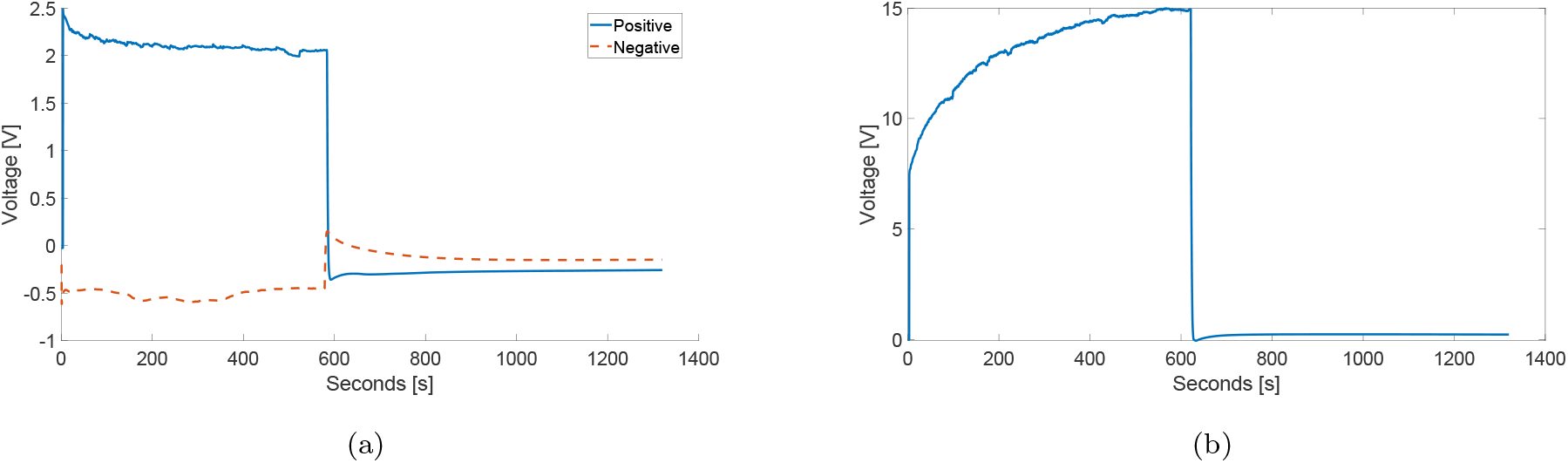
Charging mycelium to 50 V with source electrodes 10 mm apart. Substance was charged for approx. 10 minutes, readings are taken for approx. 22 minuets in total. (a) Sense electrodes were placed in series, with the source terminals (10 mm away from either positive or negative electrodes). (b) Sense electrodes were in parallel from the source electrodes (10 mm clearance). Data are discrete. Line is for eye guidance only.

**Figure 5:**
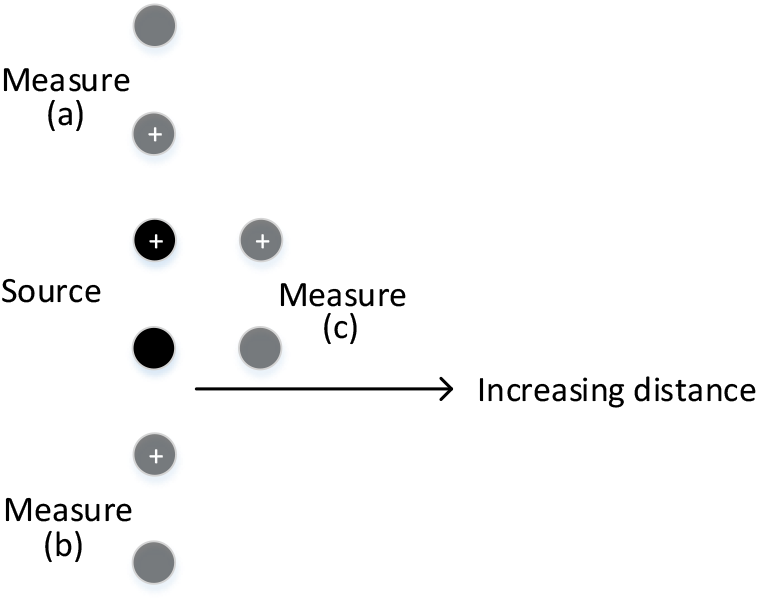
Measurement probes are arranged around the source probes to examine the charge field in the substrate.

In order to provide better contact and a more consistent data set, another batch of freshly inoculated substrate was placed in a box with gold-plated silver electrodes (multiply twisted loop, ca 1.5 cm long) with spacing of 2 cm. Changes in electrical properties of growing mycelium samples can be seen using cyclic voltammetry technique as shown in Figure 6a. These changes, however, does not seem fully systematic and the first (and only one clearly defined) jumps randomly between 0.23 and 0.36 V. This behaviour may be associated with accumulation of more redox active metabolite and indicate that simple voltammetry may not be the best technique fo monitoring of growth and evolving properties of mycelium.

**Figure 6:**
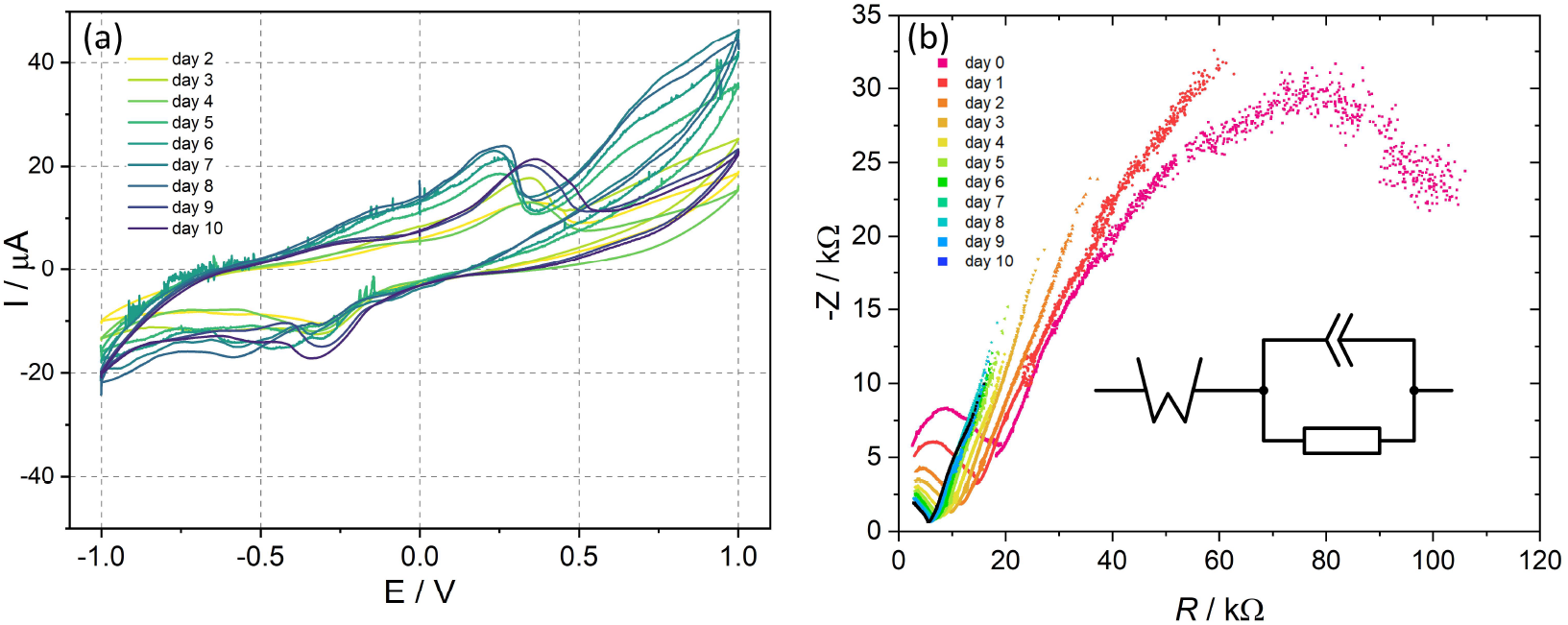
(a) Cyclic voltammograms recorded during 10 days of mycelium growth. (b) Impedance spectra recorded within 10 days after inoculation of fresh wet substrate with Pleureotus mycelium. Electrode spacing is 20 mm. An inset shown the equivalent circuit used for parameter calculation. Please note that non-ideal constant phase element is used instead of capacitor.

Impedance spectroscopy (Figure 6b) indicates a gradual changes in the spectra, reflecting the mycelium growth and demonstrating that almost fresh (wet substrate) and substrate with well-grown mycelium have completely different electrical properties (Figure 6). Observed data indicate that both capacity and resistance of the sample change gradually. In first days after inoculation changes are fast, however after day 4 changes are much smaller. Character of recorded impedance spectra indicates a complex electrical behaviour of studied samples — along with capacitive-like behaviour a significant contribution from diffusion-related Warburg-type component can be seen, especially with more matured mycelium. Therefore an equivalent circuit encompassing all these element was suggested.

The changes in the electrical properties of growing mycelium are logical and consistent with expected mycelium topology of percolated network, with percolation increasing with time (Figure 7). It can be clearly seen as gradually decreasing resistivity of the network, both Ohmic and Warburg components. Interestingly, capacitance of the network stabilised after ca. 4 days, this is also associated with decreasing capacitor quality factor. This is fully justified, as increased percolation of the network created more conductivity pathways of different time constants - proton and ionic conductivity both along the mycelium hyphae and between them. Capacitance can be associated both with electrode/mycelium interface, and contacts between different hyphae. Time course of observed changes suggests the latter as the principal component of observed capacitive behaviour.

**Figure 7:**
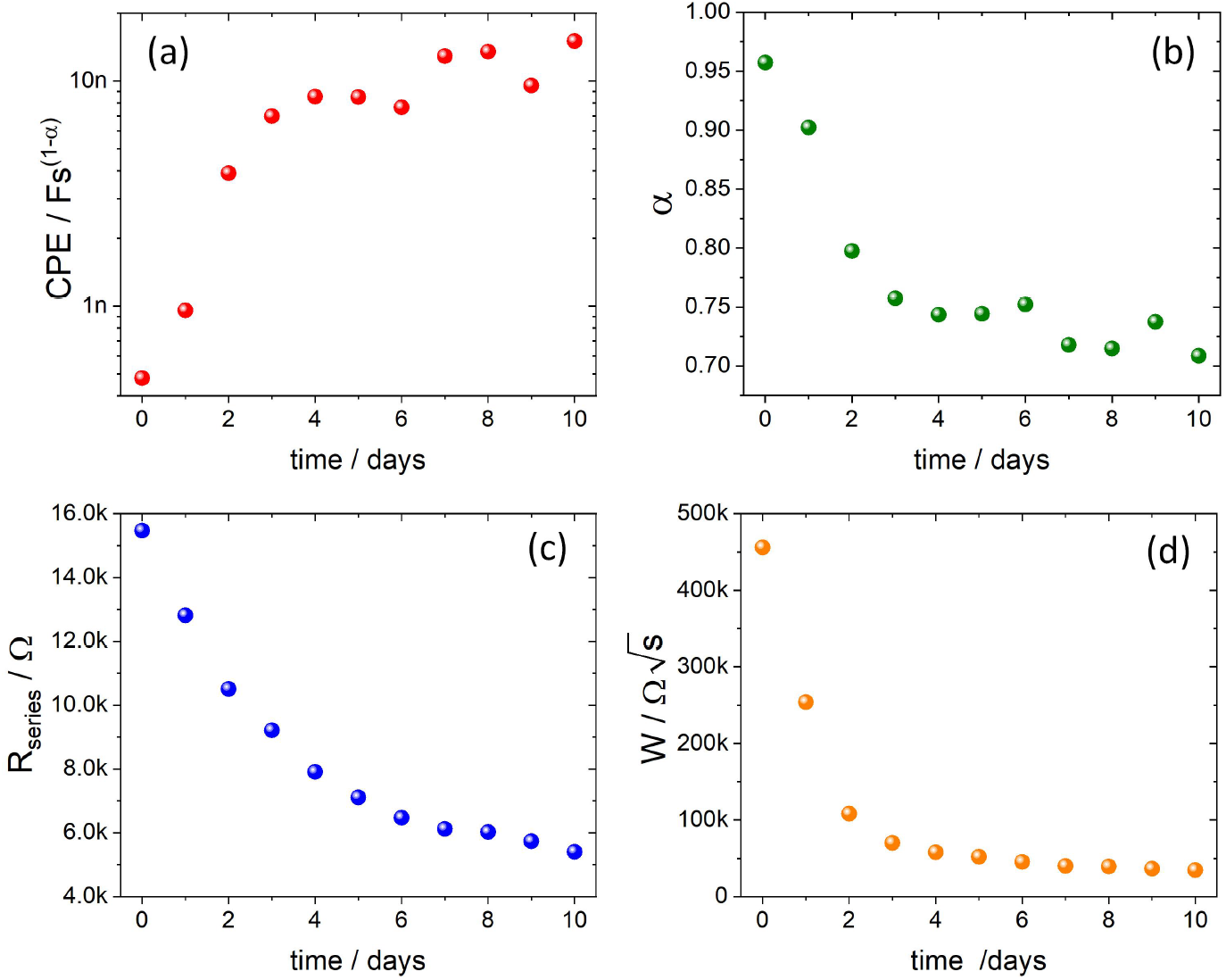
Time-dependence of the fitting parameters of the equivalent circuit from Figure 6b.

This description is a bit simplistic and does not include the whole complexity of electrical properties of mycelium. 3D representation of impedance spectra indicates, much larger complexity of their capacitive response (Fig. 8).

**Figure 8:**
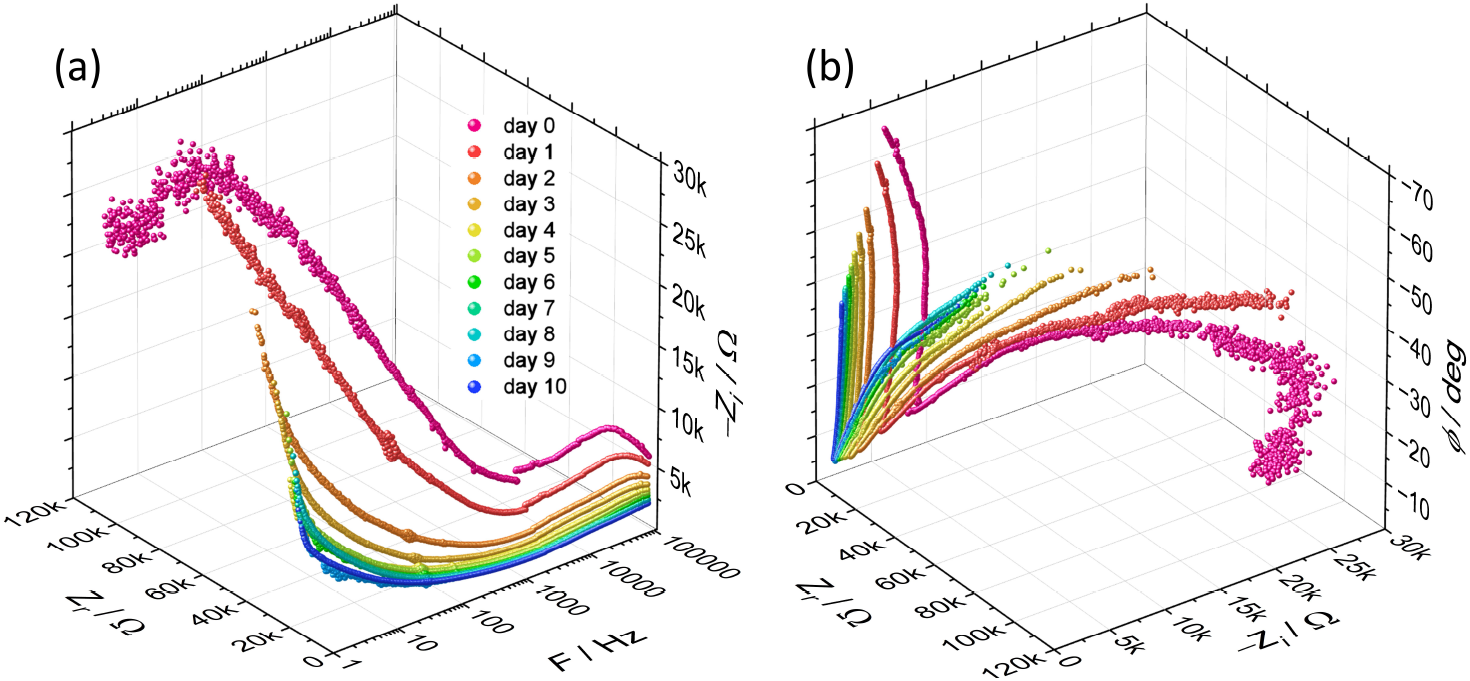
3D representation of Impedance data: (a) frequency-dependent Nyquis plot and (b) the relation between real impedance (*Z*_*r*_), imaginary impedance (*Z*_*i*_) and phase shift angle (*ϕ*).

### Charge characteristics

The charge curves in the specimens around the supply electrodes provide an insight into the ability of the substrate to conduct current. The sense electrodes were distanced from the source electrodes by varying amounts. Initially, the growth medium was studied on its own (Fig. 9). The dry wood shavings (Fig. 9(a)) showed very low voltage across the sense electrodes placed 10 mm away from the source electrodes (parallel). The dry shavings essentially acted as an open circuit and the measurement electrodes picked up noise. Damp wood shavings form a more contiguous mass and the introduction of the water helped to conduct current (Fig. 9(b)). From a 50 V source a maximum voltage of approx. 15 V was reached across the measurement electrodes. Figure 11 shows all charge curves for growth medium and mycelium. From the superimposed plots we are able to observe the maximum voltage recorded from the samples as the distance of the measurement probes are increased away from the charge probes.

**Figure 9:**
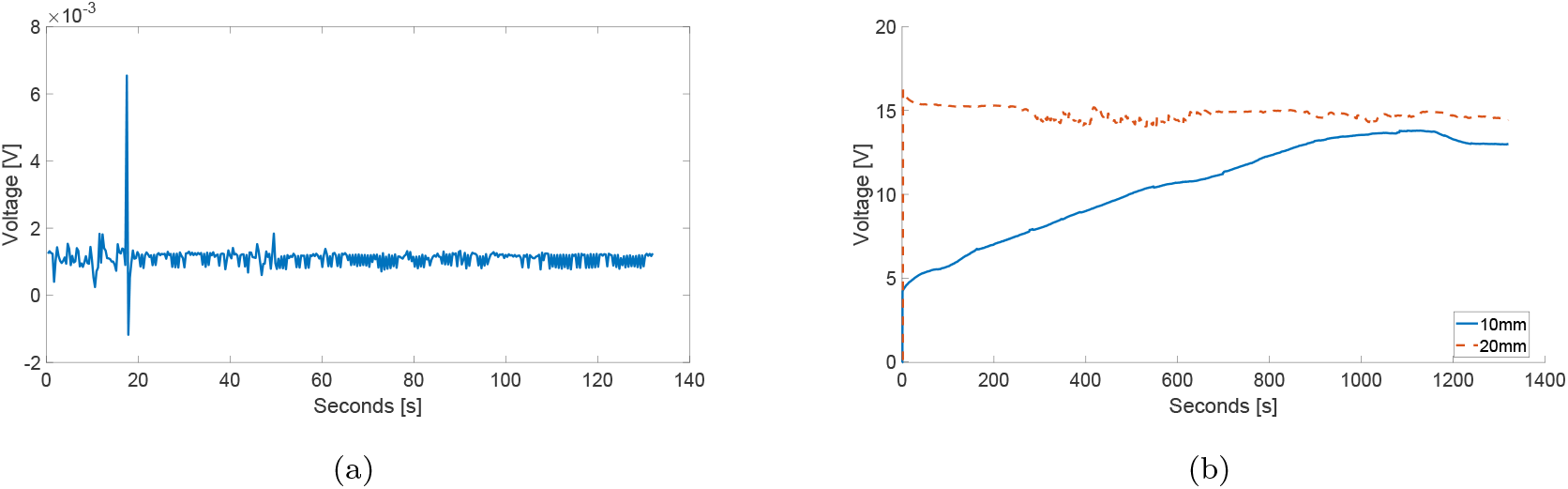
Charge dynamics of growth substrate with measurement equipment set in parallel with charge electrodes. Electrode pairs were 10 mm apart. (a) Dry wood shavings. (b) Damp wood shavings.

Preforming similar experiments with the mycelium substrate (Fig. 10), it was observed that moving the measurement electrodes further from the supply reduced the measured *V*_max_ over the 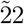 min measurement window. Figure 12 shows that the maximum measured voltage dropped rapidly as the distance from the source electrodes increased. At 10 mm away from the source, less than 1*/*5^*th*^ of the supply was measured over 22 mins, decreasing to less than 1*/*10^*th*^ at 15 mm separation. Figure 13 shows that, for the shortest paths between the two source electrodes, there was a higher current density, indicated by the field lines being closer together. As we move further away from the centre of the two probes, the current density decreased (arrows are shown further apart). Beyond the two probes in the ‘y–direction’, we would expect to experience very little current flow, however there are still fringing field effects which give rise to the small voltages shown in Fig. 4(a). Modern parallel plate capacitors have a large electrode/dielectric surface area to increase the capacitance, and multi-layer capacitors interdigitate the electrode and dielectric to achieve the required surface area in a very small package. Here we have not explored the effect of surface area on capacitance, however, it can be seen from equation (2) that increasing the area of the electrodes, in contact with our mycelium dielectric, would lead to an increase in the capacitance. This was addressed by monitoring the mycelium growth on electrodes implanted into freshly inoculated medium (*vide supra*).

**Figure 10:**
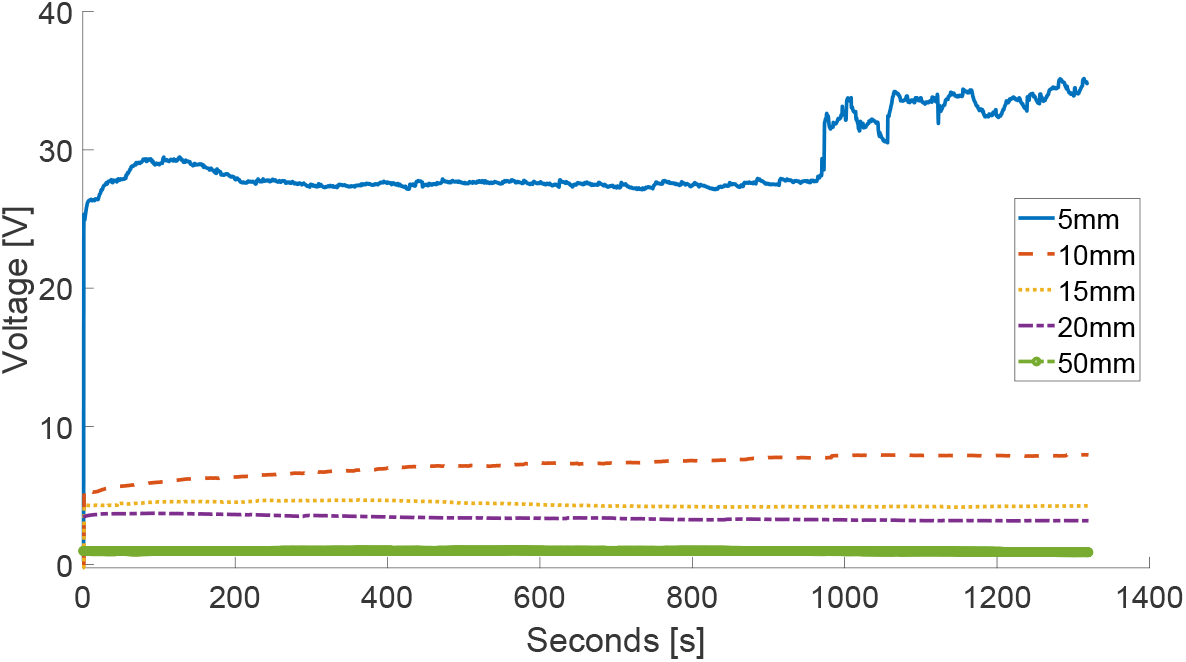
Charging of mycelium sample with measurement equipment set to measure at different distances from supply (5 mm, 10 mm, 15 mm, 20 mm, and 50 mm). Supply electrodes and measurement electrodes were arranged in parallel with each other.

**Figure 11:**
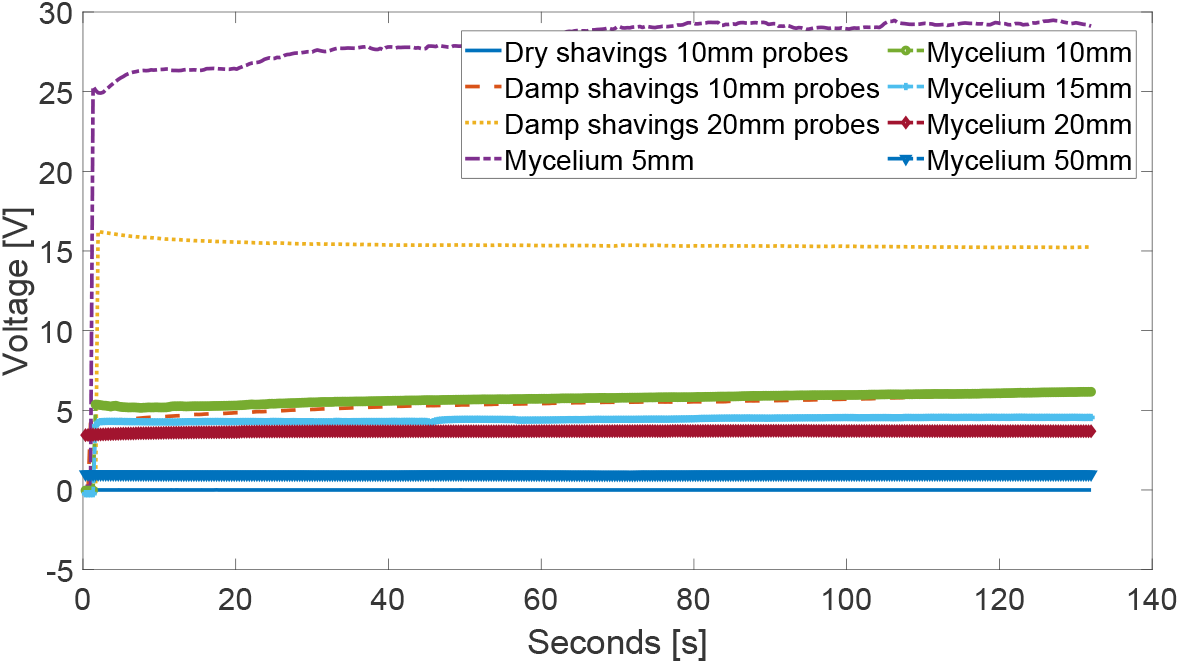
Charge characteristics of wet and dry growth medium and and mycelium with measurement probes at different distances from source probes.

**Figure 12:**
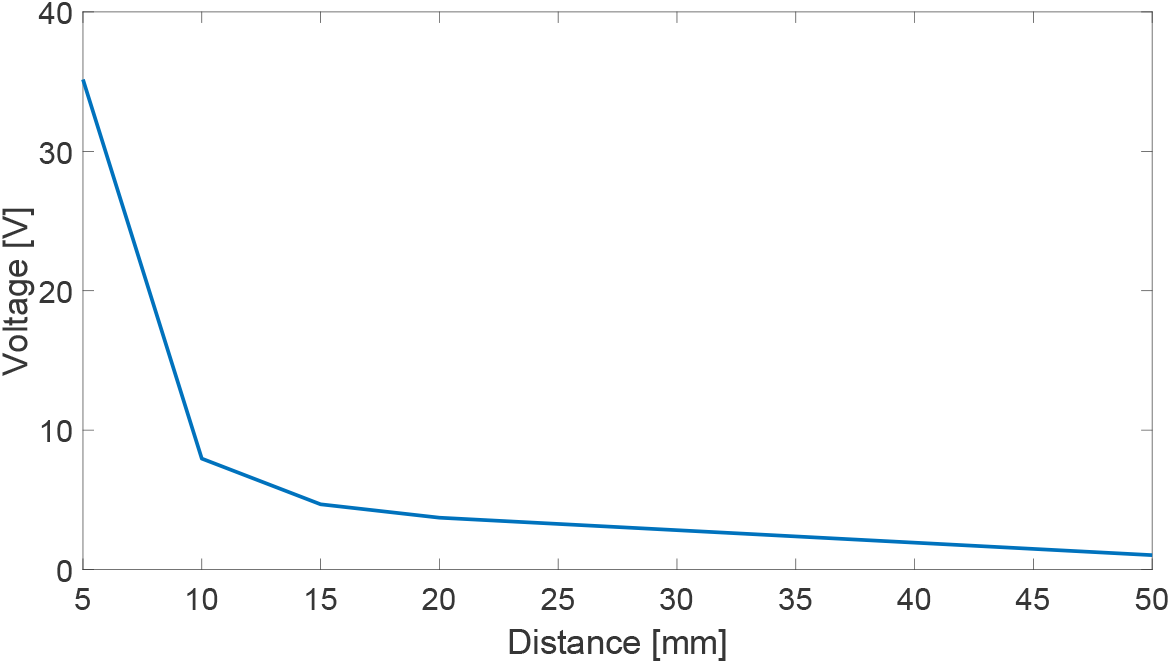
Maximum measured voltage at increasing distance from supply probes in mycelium. Data are discrete. Line is for eye guidance only.

**Figure 13:**
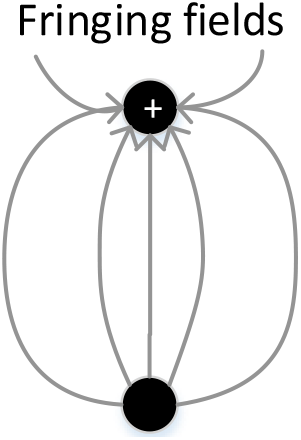
Current density between the two source electrodes. Current flow is shown in the physical direction rather than convention.

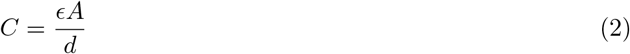

Where *C* is the capacitance, is the relative permittivity, *A* is the surface area, and *d* is the separation of the electrodes.

Additionally, the moisture content of the mycelium had an impact on its ability to conduct current. Figure. 14 shows that, if the sample of mycelium is continually charged over a period where it is also drying out, the conducted charge can decrease rapidly as the water content vanishes. The measurement electrodes for this sample are 5 mm away from the source, however we see a decrease in measured voltage of almost 15 V over the measurement window.

**Figure 14:**
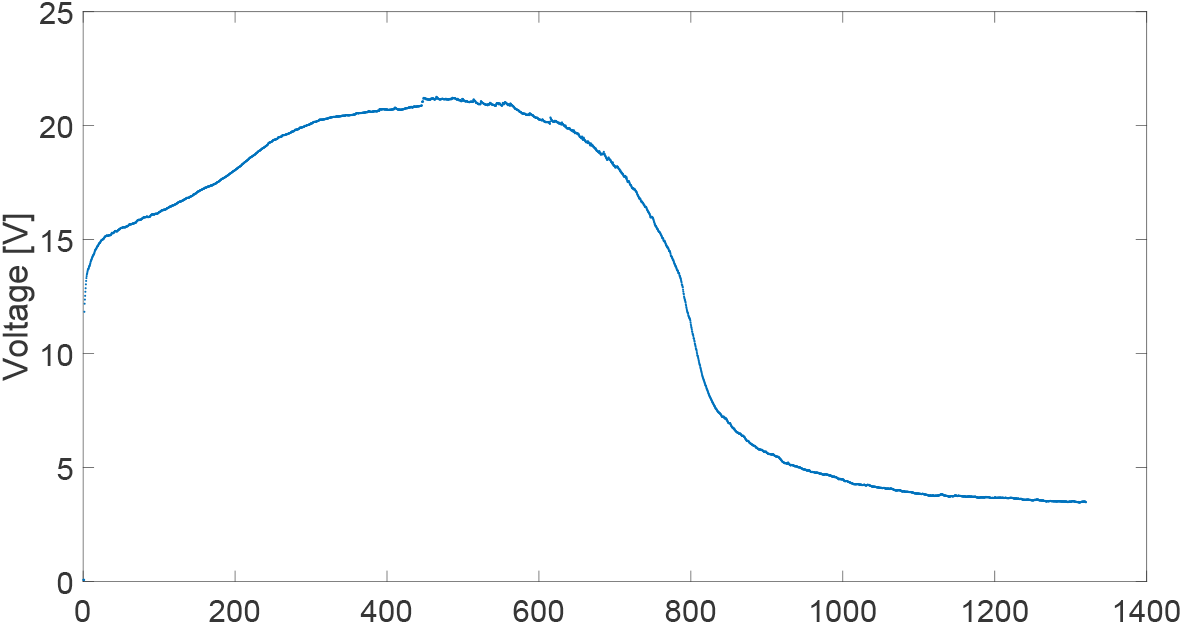
Mycelium charge measured 5 mm away from 50 V source while mycelium sample starts to dry out. Data are discrete. Line is for eye guidance only.

Application of high potentials (50 V in the case of former experiments) may, however result in electrolysis of mycelial fluid and some irreversible changes. In order to evaluate utility of fungal capacitor charge-discharge tests have been performed with much lower voltages and chronoamperimetric detection. Charging pulses with amplitudes varying from 0.2 to 1.0 V were applied to pairs of electrodes with 2 cm clearance for 30 s, and then discharge current at short circuit conditions was recorded for additional 90 s. Resulted current profiles are shown in Figure 15a. It can be observed, that effective total capacitance (Figure 15b) is much higher than results obtained at low amplitude impedance spectroscopy (Figure 7). It can be justified by a mixed capacitance character with a dominating contribution of electrochemical pseudocapacitance, additionally with a significant Faradaic component (please note quasi-reversible peaks in Figure 6a. We can hypothesise, that ionic processes at the surface of biological membranes are the dominating factor, as the voltage dependence of capacitance is monotonous.] within experimental error. This excluded Faradaic component as a dominating one, however its importance should be also noted. It is further supported by observed discrepancy between charge stored during charging pulse and charge collected during discharge. This discrepancy is voltage-independent and indicates significant charge loss, most probably due to ionic diffusion within hyphae.

**Figure 15:**
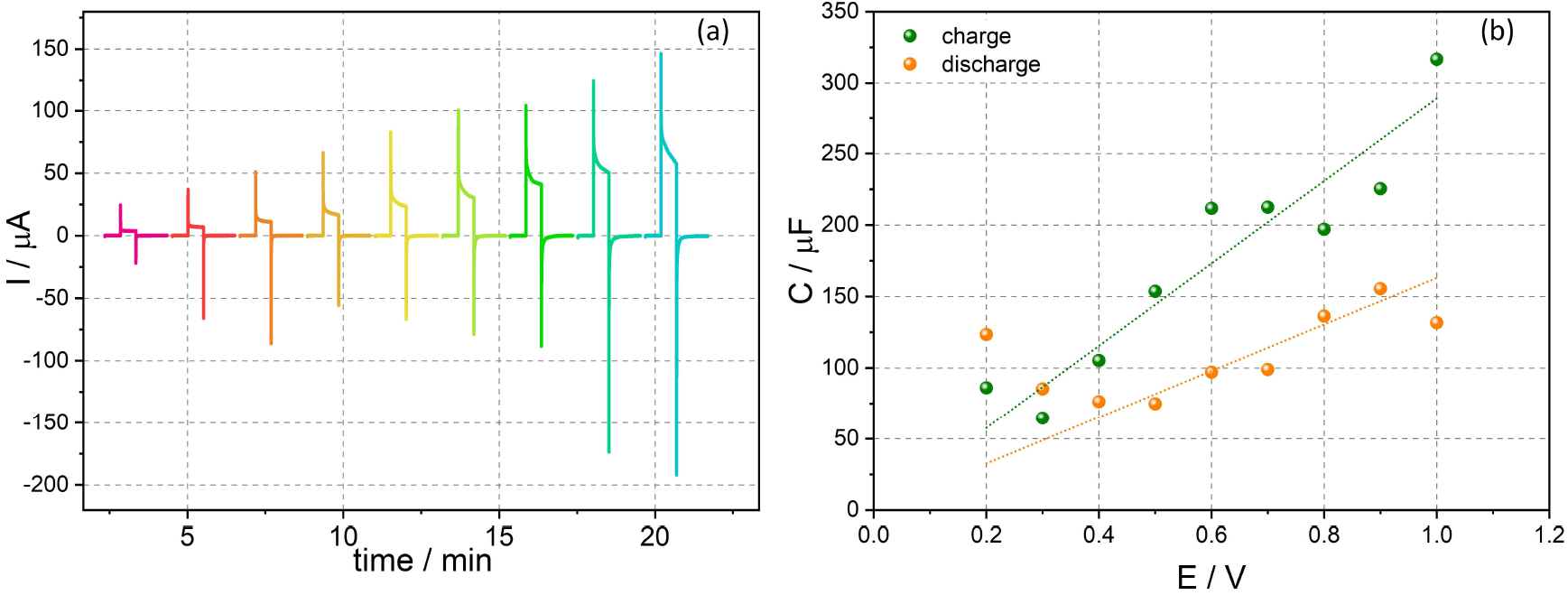
(a) Charge/discharge current profiles. (b) Capacitance vs applied voltage for calculated by integration from appropriate parts of the charge/discharge curves.

## 4 Conclusions

Mycelium exhibits rather unconventional, voltage- and frequencydependent capacitive characteristics. Depending on the frequency measurement and applied voltage, capacitance of mycelium samples very form hundreds of picofarads to hundreds of microfarads. This enormous discrepancy, spanning six orders of magnitude, is, however not unjustified. In reflects complexity of electrical behaviour of mycelium, which is a consequence of its molecular composition, structure and topology. We cannot expect simple electrical behaviour from wet fibrous hemp-derived substrate, overgrown with living mycelium. Complex topology of the materials, combined with biochemical processes creates a challenging system to study. Simple measurements with benchtop multimeter do not yield high capacitance values, probably due to non-optimal contact of electrodes with mycelium and substrate. High voltage measurements, it turn, may induce some irreversible processes and electrolysis of mycelial cytosol, followed by cell disruption. Not harmful to the mycelium as a whole, these local damages are reflected in rather low charge storage capability due to damaged cell walls. AC measurements also do not give high capacintance values (up to 10 nF), suggesting contribution of diffusive processes (observed as Warburg impedance in impedance experiments) Finally, at low applied voltages, DC experiments demonstrate, however, high charge storage capacity significantly increased up to 150 microfarads. Due to leackage processes and rather low quality factor they cannot be efficiently used for energy storing application, however may be further explored for other application, e.g. sensing, mycelial communication and monitoring of mycelium growth. This shows great potential for the use of mycelium networks to conduct or store charge in local ‘hot spots’ that are isolated from other areas in the immediate vicinity, especially at low signal amplitudes. However, it is crucial that the moisture content of the mycelium is kept constant since the ability to carry charge is strongly influenced by moisture content.

Any potential analogue circuits implemented with live mycelium will be vulnerable to environmental conditions, especially humidity, availability of nutrients and removal of metabolites. Ideally, the mycelium networks should be stabilised so they continue functioning whilst drying. This stabilisation can be achieved either by coating or priming the mycelium with polyaniline (PANI) or poly(3,4-ethylenedioxythiophene) and polystyrene sulfonate (PEDOT-PSS). This approach has been proven to be successful in experiments with slime mould *P. polycephaum* [49, 49, 50], and thus it is likely that a similar technique may be applied to fungi. Moreover, PABI and PEDOT-PSS incorporated in, or interfaced with, mycelium can bring additional functionality in terms of conductive pathways [51], memory switches [52, 53] and synaptic-like learning [54, 55]. An optional route toward the functional fixation of mycelium would be doping the networks with substances that affect the electrical properties of mycelium, such as carbon nanotubes, graphene oxide, aluminium oxide, calcium phosphate. Similar studies conducted in our laboratory using slime mould and plants have shown that such an approach is feasible [56, 57]. Moreover using a combination of PANI and carbon nanotubes in the mycelium network afford it supercapacitive properties [58, 59]. Another potential direction of future studies would be to increase the capacity of the mycelium as a result of modifying the network geometry by varying nutritional conditions and temperature [60, 61, 62, 63], concentration of nutrients [64] or with chemical and physical stimuli [41]. With regards to the impact of our finding for the field of unconventional computing, we believe further research on experimental laboratory implementation of capacitive threshold logic [65, 66], adiabatic capacitive logic [67] and capacitive neuromorphic architectures [68] will yield fruitful insights.

## Acknowledgements

Authors thank professor Kapela Pilaka for numerous valuable comments and fruitful discussions. This project has received funding from the European Union’s Horizon 2020 research and innovation programme FET OPEN “Challenging current thinking” under grant agreement No 858132. KM and KS acknowledge the support from the programme “Excellence initiative–research university” for the AGH University of Science and Technology.

## Author contributions

A.B. and A.A. conceived the idea of experiments. A.A. and A.P. prepared the substrate colonised by mycelium. A.B. and K.S. performed experiments, collected data and produced plots in the manuscript. Electrochemical data were analysed by KM, who performed also an equivalent circuit modelling. A.B. and A.A. prepared manuscript (wrote and reviewed all contents). A.P. reviewed manuscript.

